# Heat Shock in *C. elegans* Induces Downstream of Gene Transcription andAccumulation of Double-Stranded RNA

**DOI:** 10.1101/448233

**Authors:** Marko Melnick, Patrick Gonzales, Joseph Cabral, Mary A. Allen, Robin D. Dowell, Christopher D. Link

## Abstract

We observed that heat shock of *Caenorhabditis elegans* leads to the formation of nuclear double-stranded RNA (dsRNA) foci, detectable with a dsRNA-specific monoclonal antibody. These foci significantly overlap with nuclear HSF-1 granules. To investigate the molecular mechanism(s) underlying dsRNA foci formation, we used RNA-seq to globally characterize total RNA and immunoprecipitated dsRNA from control and heat shocked worms. We find antisense transcripts are generally increased after heat shock, and a subset of both sense and antisense transcripts enriched in the dsRNA pool by heat shock overlap with dsRNA transcripts enriched by deletion of *tdp-1,* which encodes the *C. elegans* ortholog of TDP-43. Interestingly, transcripts involved in translation are over-represented in the dsRNAs induced by either heat shock or deletion of *tdp-1.* Also enriched in the dsRNA transcripts are sequences downstream of annotated genes (DoGs), which we globally quantified with a new algorithm. To validate these observations, we used fluorescence *in situ* hybridization (FISH) to confirm both antisense and downstream of gene transcription for *eif-3.B,* one of the affected loci we identified.

## Introduction

Cytoplasmic proteotoxic stress induced by temperatures outside of the optimal range for cells or organisms triggers the heat shock response (HSR) [1]. The response to heat shock is multi-faceted and regulation of both transcription and translation occurs. Transcriptional responses include formation of stress granules, alternative splicing, and aberrant transcriptional termination [2–5]. The HSR is a highly conserved transcriptional response and is driven largely by the heat shock transcription factor HSF1 [6]. Under basal level conditions, HSF1 is a monomer in the cytoplasm and nucleus. Upon stress, HSF1 undergoes homotrimerization and binds to DNA heat shock elements (HSE) and initiates the transcription of heat shock protein genes [7, 8]. In addition, translation of non-heat shock mRNAs is reduced through pausing of translation elongation as well as inhibition of translation initiation [9–11]. Regulation and clearance of misfolded proteins by heat shock proteins has been implicated in neurodegenerative diseases such as Huntington’s disease (HD), Parkinson’s disease (PD), Alzheimer’s disease (AD), and amyotrophic lateral sclerosis (ALS) [12].

Aside from the canonical binding of HSF1 to HSE loci, heat shock can cause HSE-independent transcriptional changes [2]. In mammalian cells, HSF1 granules co-localize with markers of active transcription where HSF1 binds at satellite II and III repeat regions [13]. In the worm *Caenorhabditis elegans,* HSF-1 granules also show markers of active transcription but the putative sites of HSF-1 stress granule binding are unknown [14].

In addition to formation of HSF1 stress granules, heat shock can cause reduced efficiency of transcription termination and the accumulation of normally untranscribed sequences, designated in the literature as downstream of gene-containing transcripts (DoGs) [5]. In eukaryote transcriptional termination, the Carboxyl-terminal domain (CTD) of RNA Polymerase II (Pol II) interacts with a complex of cleavage and polyadenylation (CPA) factors responsible for generating the polyadenylate [Poly(A)] tail at mRNA 3’ ends. Two models exist for how the pre-mRNA poly(A) site (PAS) contributes to transcription termination. The allosteric model proposes that Pol II senses PAS during elongation leading to a conformational change in the Pol II active site eventually leading to Pol II release. The other, dubbed the torpedo model, proposes that the nuclear 5’-3’ exonuclease Xrn2 is recruited to the PAS and triggers Pol II release when it degrades the downstream transcript and catches up to elongating Pol II [15].

Recent studies have shown increased antisense transcription when DoGs/Read-through transcription goes past the PAS into neighboring genes on opposite strands [16–19]. Antisense transcription has the potential to modulate gene expression by creation of double-stranded RNA (dsRNA) with subsequent degradation through RNA interference (RNAi) [20].

Previous studies in our lab found deletion of *tdp-1*, the worm ortholog of ALS associated protein TDP-43, results in the accumulation of dsRNA foci [21]. In addition to deletion of *tdp-1*, we discovered that heat shock robustly induced nuclear dsRNA foci in worms. To assay this unexpected formation of dsRNA, we performed strand-specific RNA-seq and strand-specific RNA-immunoprecipitation sequencing (RIP-seq) with the J2 antibody specific for dsRNA. In heat shocked worms, we find increased J2 enrichment of downstream-of-gene transcripts as well as genes involved in translation. To identify altered transcription genome-wide, we developed an algorithm called Dogcatcher that provides DoG locations, differential expression of DoGs, and genes that overlap with DoGs on the same or opposite strand.

## Materials and Methods

### *Caenorhabditis elegans* culturing and strains

Hermaphrodites from each strain were kept at 16 °C on Nematode Growth Media (NGM) plates seeded with Escherichia coli strain OP50 as a food source according to standard practices [22]. To obtain age synchronized worms, we used alkaline hypochlorite bleach on gravid adults to obtain eggs that were hatched overnight in S-basal buffer [23]. Worms were then allowed to grow to 1-day-old adults (approximately 80h at 16 °C). List of strains used in this study is available in S1 Table.

### Heat stress treatment

Heat stress treatment was performed in an air incubator set to 35 °C for 3 hours for the RNA-seq experiments. After stress, populations were washed off with S-basal buffer and immediately fixed for immunohistochemistry and/or fluorescence in situ hybridization (FISH), flash frozen in liquid nitrogen for quantitative reverse transcriptase polymerase chain reaction (qRT-PCR), or crude extracts were created with subsequent J2-Immunoprecipitation (J2-IP) as previously described [21].

### RNA isolation, cDNA library preparation, and RNA Sequencing

Total RNA was extracted from worms using TRIzol (Invitrogen #15596026) extraction. Chloroform was used to solubilize proteins and TURBO DNase (Invitrogen) was used to remove DNA. For total RNA libraries, 5 μg of RNA was ran through a RiboZero column (Epicenter, #R2C1046) to remove ribosomal RNA. Libraries were created using Illumina TruSeq kits (RS-122-2001). RNA recovered by immunoprecipitation with the J2 antibody of young adult worms as well as input material (as a loading control) was converted into strand-specific total RNA libraries using V2 Scriptseq (Epicenter #SSV21106) kits following manufacturer’s instructions, except reverse transcription was done with SuperScript III (Invitrogen #18080 044) using incrementally increasing temperatures from 42 to 59 °C to allow for transcription though structured RNAs. rRNA was not removed from J2-IP RNA samples. Libraries were sequenced on an Illumina HiSeq 2000 platform at the Genomics Core at the University of Colorado, Denver. Data were deposited under GEO accession number GSE120949.

### Immunohistochemistry and Fluorescence in situ Hybridization (FISH)

For immunohistochemistry, all washes used a constant volume of 1ml sterile S-basal buffer unless otherwise noted. Worms were first washed off plates, spun down into a pellet, and fixed in 4% paraformaldehyde. Worms were then resuspended in 1ml of Tris/Triton buffer with 5% beta-mercaptoethanol and incubated in a rocker for two days at 37°C. After two days, worms were washed two times and put into collagenase buffer. Next, worms were placed into a 1:1 dilution of 1mg/ml type IV collagenase (Sigma) and S-basal buffer for 45 minutes at 37 °C with rocking. Worms were checked under the microscope to ensure cuticle breakage then quenched in cold Antibody buffer A (1X Phosphate buffered saline, 0.1% Bovine Serum Albumin, 0.5% Triton X-100, 0.05% Sodium Azide). Worms were then washed, pelleted, and primary antibodies were added for 16 hours at 4°C. Next, worms were washed twice in Antibody buffer B (same as Antibody buffer A except using 1% Bovine Serum Albumin), pelleted, and secondary antibodies were added with subsequent incubation for 2 hours at room temperature. Finally, worms were washed twice in Antibody buffer B and then placed in 50/mul of Antibody buffer A. Permeabilized worms were probed with the primary J2 antibody (English and Scientific Consulting Lot: J2-1102 and J2-1103) at 4qμg/mL and secondary antibody Alexa dye-conjugated goat anti-mouse at 4μg/mL. DAPI nuclear stain was added along with secondary antibodies at 5μg/mL to visualize nuclei.

Stellaris FISH probes (Biosearch technologies) [24] were custom designed using the Stellaris RNA FISH probe designer. Three regions were chosen for probing, and each probe was tested against the *C. elegans* genome using BLAST to identify any complementarity to nontarget sequences. A probe was excluded if it was in an intron, had a highly repetitive sequence outside of the region, or matched other regions up to 18nt long with high transcriptomic expression viewed in the Integrative Genome Viewer (IGV) [25] Probes and locations are available in S1 File.

For FISH probing and storage, the Stellaris protocol for *C. elegans* was followed using RNAase OUT (Invitrogen) when applicable. Briefly, worms were washed off plates using nuclease-free water and fixed for 45 minutes at room temperature in a fixation buffer (1:1:8 of 37% Formaldehyde, 10X RNase-free Phosphate Buffered Saline (PBS), Nuclease-free water). Worms were then washed twice with 1X RNAase-free PBS and permeabilized in 70% ethanol overnight at 4°C. Worms were then incubated at room temperature in Stellaris Wash Buffer A, pelleted, and incubated for 16 hours in a 37 °C water bath in the dark with 100μl of the probe/hybridization buffer (9:1 of μl Stellaris RNA FISH Hybridization buffer, Deionized Formamide with a 100:1 Hybridization buffer, FISH probe). Next, 1mL of Stellaris Wash Buffer A was added with 30 more minutes of incubation in the dark 37 °C water bath. Stellaris Wash Buffer A was then aspirated and incubated with DAPI (1:1000 of 5μg/m DAPI, Stellaris Wash Buffer A) for 30 more minutes of the dark 37 °C water bath. Lastly, the DAPI buffer was aspirated and 1mL of Stellaris Wash Buffer B was added with a 5 minutes room temperature incubation.

A modification of the immunohistochemistry protocol was used when doing immunohistochemistry and FISH. The immunohistochemistry protocol was the same except all washes were done using RNAase-free PBS or water and RNAase-free reagents (Tris/Triton Buffer, Collagenase Buffer, Collagenase, Antibody Buffer A, Antibody Buffer B) were created by adding RNAase OUT (2:10000 of RNAase OUT, reagent). After antibody staining, the FISH protocol was started at the probe/hybridization step.

## Microscopy

Images were acquired with a Zeiss Axiophot microscope equipped with digital deconvolution optics (Intelligent Imaging Innovations). Image brightness and contrast were digitally adjusted in Photoshop.

### Quantification of occurrence of HSF-1 and J2 foci over time

Intestinal nuclei of the worms were isolated from the rest of the image and the Foci Picker3D plug-in was used to count foci. The FITC channel of the image was converted to 16 bit and analyzed. Foci Picker3D settings were changed from default by changing the Minlsetting to 0.25 and the ToleranceSetting to 20 before running analysis. Five worms were selected for each time point. Raw data is available in S2 Table.

### Data Analysis

Detailed instructions on algorithms and analysis is provided in S3 file. Briefly, reads were checked for quality with FastQC v0.11.7 [26], adapters were trimmed using Trimmomatic-0.36 [27] (S2 file), and reads were aligned to the worm genome WS258 using STAR-2.5.2b [28]. Genes and DoGs (identified by Dogcatcher, described below) were assigned counts using Rsubread v1.28.1 featureCounts [29] and were rRNA-normalized according to the rRNA subtraction ratio (RSR) (described in supplemental). Differential expression was obtained using DESeq2 v1.20.0 and the likelihood ratio test (LRT) set with Total/Input and J2 treated as separate variables within the condition [30].

We created an algorithm called Dogcatcher to identify and analyze DoGs. Briefly, Dogcatcher uses a sliding window approach to identify contiguous regions of transcription above a defined threshold. If the sliding window runs into a gene on the same strand it will either continue (meta read-through) or stop (local read-through). Dogcatcher will output bedfiles, gtfs and dataframes of all DoGs and antisense DoGs for a sample along with differential expression and genes overlapping DoGs (For additional details see bioinformatics supplemental S3 File). For improved normalization in DESeq2, non-significant genes are added when calculating differential expression and removed afterward. The Dogcatcher algorithm and README is available at https://github.com/Senorelegans/Dogcatcher. For processing J2 enrichment, a modified version of Dogcatcher was used that applies the rRNA subtraction ratio normalization and likelihood ratio test from DESeq2 (available at https://github.com/S enorelegans/heatshock_and_tdp-1_dsRNA_scripts/J2_enrichment_Dogcatcher). After Dogcatcher was used, DoGs overlapping operons on the same strand were removed (S3 File for operon removal methods). All of the scripts used to process the data and create figures can be found at https://github.com/Dogcatcher/heatshock_and_tdp-1_dsRNA_scripts

## Results

### Heat shock induces nuclear dsRNA foci in *C. elegans*

While looking for conditions that might induce dsRNA foci besides loss of *tdp-1*, we found that heat shock robustly induced dsRNA nuclear foci. Upshifting wild type worms to 35 °C or 37 °C induced foci detectable with the J2 dsRNA-specific monoclonal antibody within 30 minutes, primarily visible in intestinal and hypodermal nuclei. To determine if these foci overlapped with previously identified nuclear HSF-1 stress granules, we repeated the heat shock experiment with strain OG497 *(drSI13)* [14]. This strain has a single copy insertion of *hsf-1* with a C-terminal GFP driven by the *hsf-1* promoter, and shows nuclear GFP expression that redistributes into granules after a one minute heat shock at 35 °C [14]. Using the J2 antibody for immunohistochemistry, we found J2 dsRNA foci in nuclear regions that partially overlapped with nuclear HSF-1 stress granules when drSI-13 worms were heat shocked for 35 °C for 40 minutes (Fig 1).

**Fig.1.**
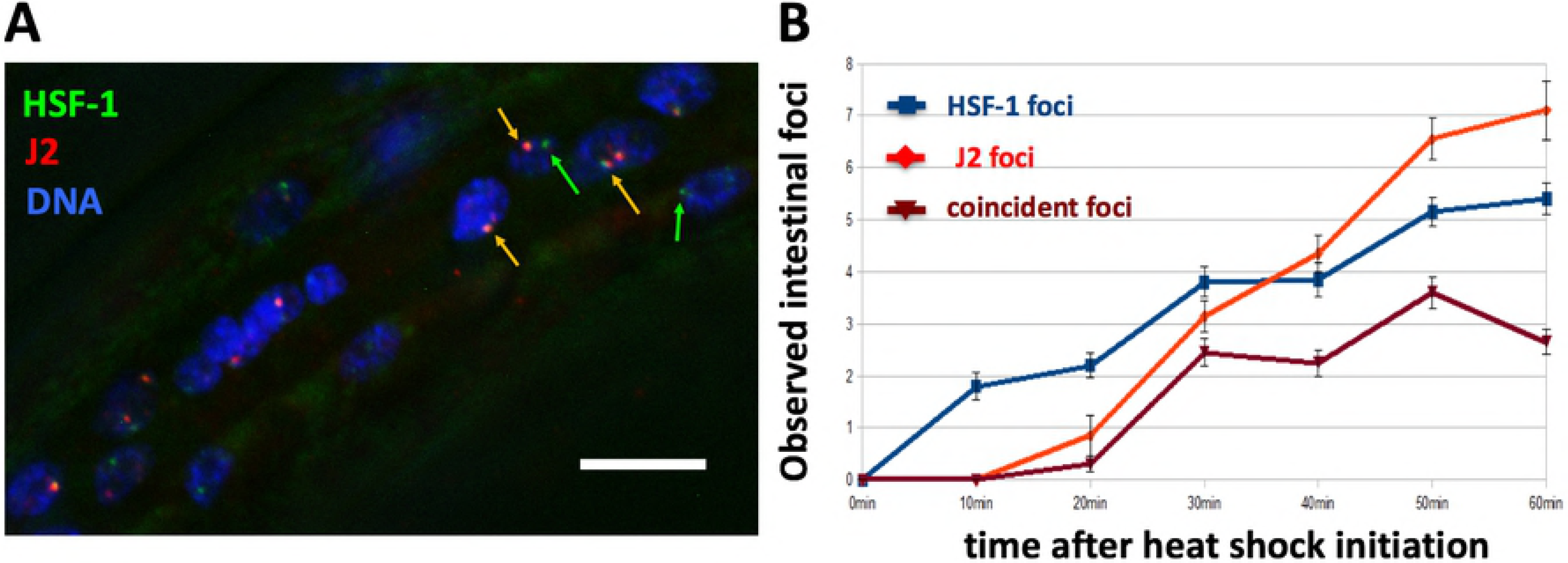
Heat shock induces nuclear foci detectable with dsRNA-specific antibody J2. (A) Mid-animal intestinal region of 4th larval stage drSI13 worm fixed 40 minutes after heat shock at 35° C. Note nuclear J2 foci (orange arrows), many of which overlap with HSF-1 foci. HSF-1-only foci indicated by green arrows. White size bar in bottom right corner (20 microns across). (B) Quantification of occurrence of HSF-1 and J2 foci over time, 4 intestinal nuclei per worm scored.

### Recovery of dsRNA by J2 immunoprecipitation

In order to identify dsRNA transcripts induced by heat shock, we performed strand specific RNA sequencing (RNA-seq) and strand specific RNA immunoprecipitation sequencing (RIP-seq) (Fig 2). Total/Input RNA and RNA immunoprecipitated with the J2 antibody was extracted and sequenced for heat shocked N2 (wild type) worms (in duplicate) and non-heat shocked worms (in triplicate). The J2 antibody is specific for dsRNA 40bp or more [31]. Importantly, transcripts from the J2 IP could include full length dsRNA transcripts or single stranded RNA (ssRNA) with 40bp or more sections of dsRNA. dsRNA can occur via base pairing with a different transcript (interstrand) or self-complementarity within the same transcript (intrastrand).

**Fig.2.**
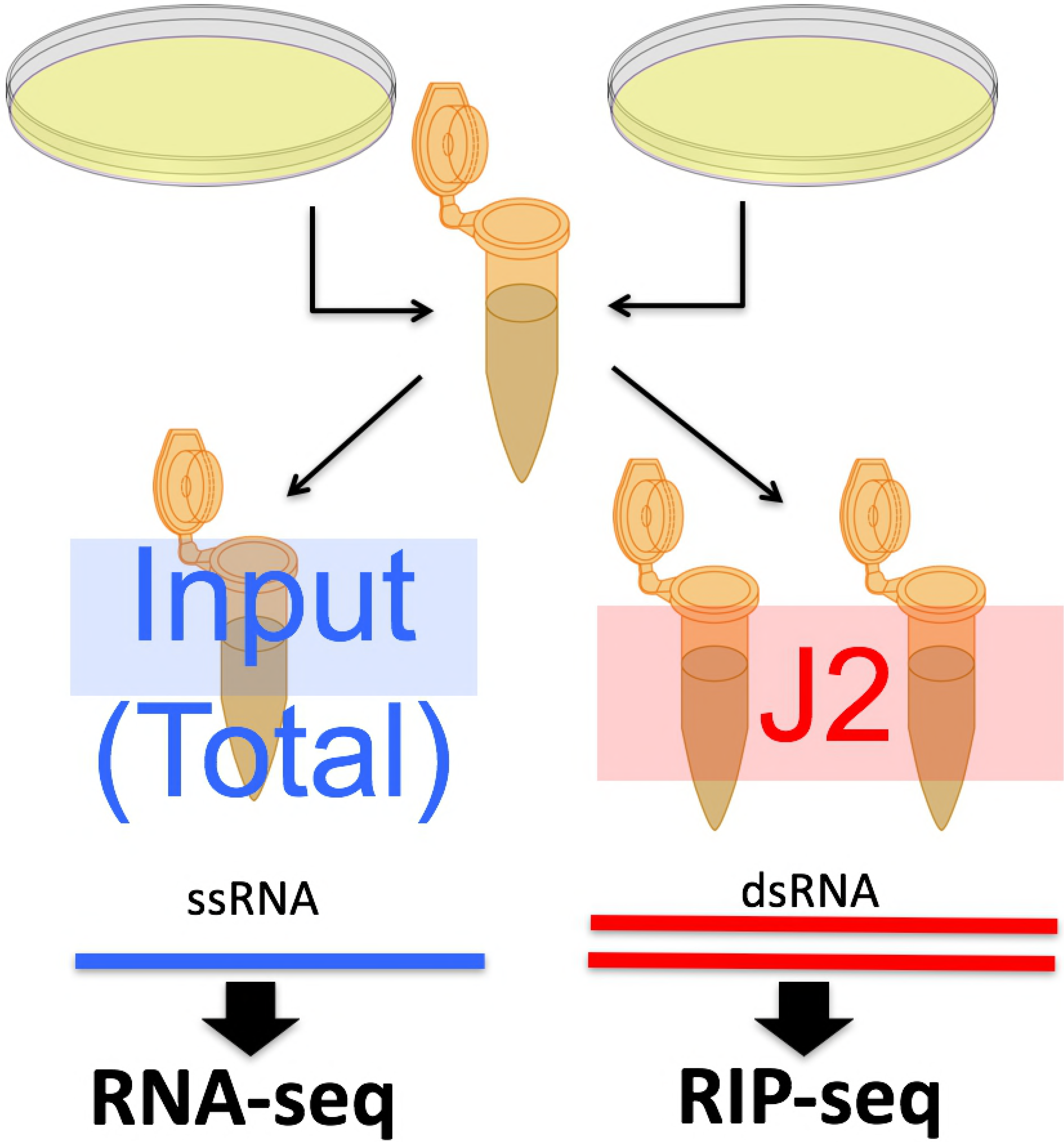
Schematic of recovery of RNA pools for high throughput sequencing analysis. Control and heat shocked worm populations were recovered and lysed. Worm lysates were then split to recover total input RNA or immunoprecipitated with the J2 antibody.

### Antisense transcripts increase after heat shock

The apparent increase in dsRNA we observed in heat shocked worms [and previously observed in the *tdp-1(ok803)* mutant] could result from an increased accumulation of antisense transcripts. To obtain a global view of antisense levels, we calculated an antisense/sense ratio for genes using the total RNA samples (S3 File for methods). Genes with a minimum of 20 antisense and 20 sense reads were used for the analysis. For both heat shock and *tdp-1(ok803)* samples, we found a strong trend towards increased antisense/sense ratios compared to normal conditions (Fig 3). Heat shock results in 77% of the genes tested (785/1016) having increased antisense/sense ratios > 1 compared to wild type, and *tdp-1(ok803)* having 86% (799/925) with increased ratios compared to wild type. Notably, heat shock produced a higher average antisense/sense ratio with a greater spread of ratios compared to *tdp-1(ok803).*

**Fig.3.**
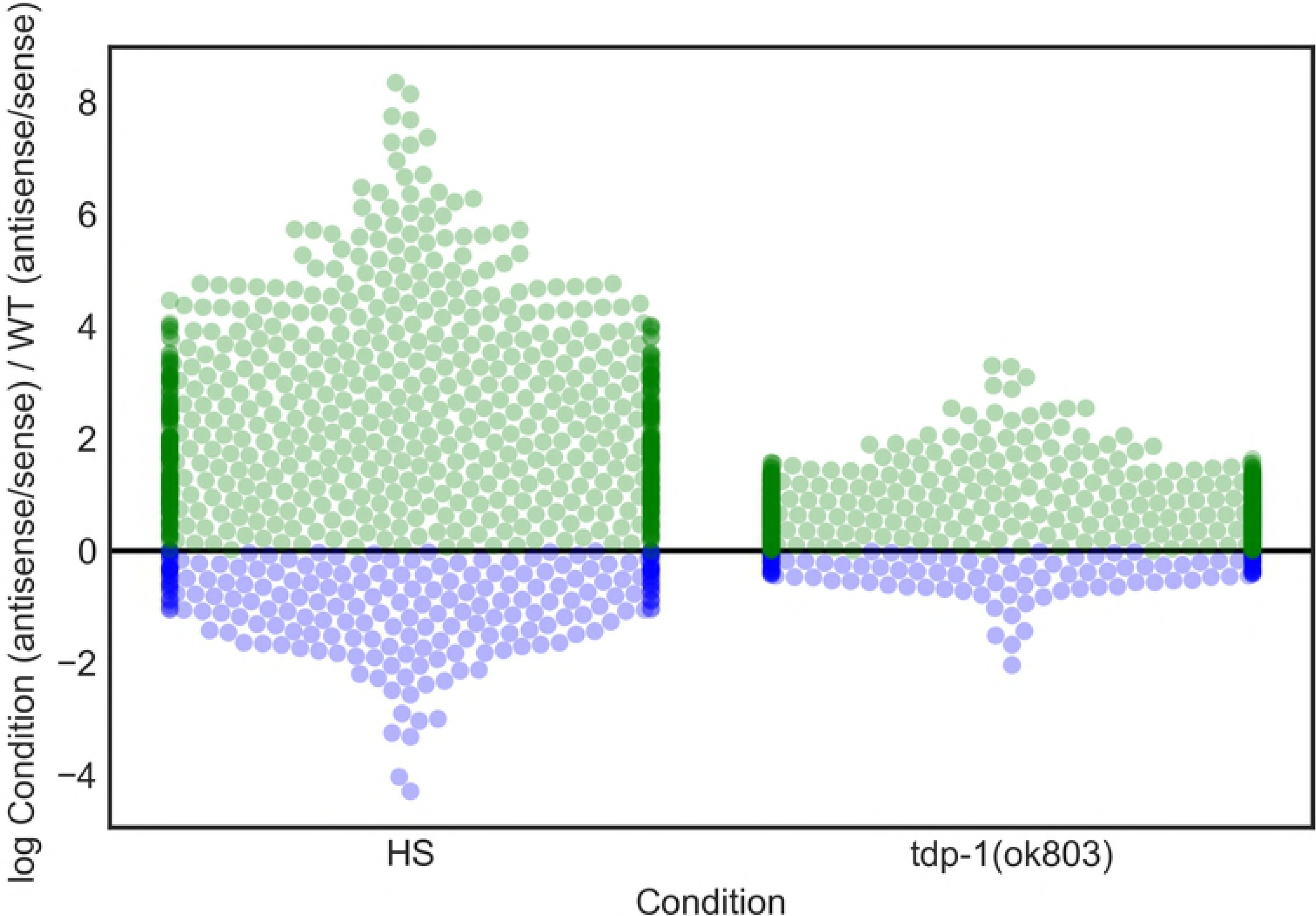
Quantification of genes with changes in antisense/sense rations after heat shock or deletion of *tdp-1* in total RNA. 785 genes up in antisense and 231 genes down in antisense for heat shock (HS) compared to wild type (WT) using the ratio of antisense/sense. 799 genes up in antisense and 126 genes down in antisense for *tdp-1(ok803)* using the ratio of antisense/sense. Colored dots represent genes with increased antisense (green) and decreased antisense (blue) compared to wild type (n=1).

### Comparison of dsRNAs identified in worms heat shocked or deleted for *tdp-1*

Considering that both heat shock and deletion of the *tdp-1* gene lead to the formation of nuclear dsRNA foci, we sought to determine if this phenotypic similarity also extends to transcriptional changes. After heat shock, we found that a large number of gene transcripts were significantly (FDR <0.05) enriched (4774) or depleted (1669) in the pool of RNAs immunoprecipitated by the dsRNA-specific antibody J2 (relative to untreated worms) (Fig 4A). We also identified antisense transcripts with significantly altered representation in the J2-immunoprecipitated pool, and found 650 were enriched and 477 depleted (Fig 4B). A minority of genes had both sense and antisense transcripts significantly enriched (180) or depleted (48) in the heat shock J2-IP pool (Fig 4A-B) (plot of genes showing only significant sense and antisense transcription for heat shocked worms in S1 Fig). In *tdp-1(ok803)* significant (FDR <0.05) gene transcripts, we found a smaller number of sense enriched (418) and depleted (59) (Fig 4C), as well as antisense enriched (245) and depleted (14) genes (Fig 4D). Similar to heat shock, *tdp-1(ok803)* had relatively fewer genes with both sense and antisense transcripts significantly enriched (6) and depleted (1) (Fig 4C-D) (plot of genes showing only significant sense and antisense transcription for *tdp-1(ok803)* worms in S2 Fig). We found a significant [P < 1 × e^-30^, hypergeometric (hgd)] overlap of J2-enriched gene transcripts between the heat shock and *tdp-1*(*ok803*) populations in both sense (Fig 4E) and antisense (Fig 4F), suggesting that there might be some similarities between the transcriptional changes induced by heat shock and deletion of *tdp-1.* However, with J2-depleted transcripts, we found no significant overlap in (P = 0.165, hgd) in sense transcripts and no significant overlap in antisense transcripts (P = 0.187, hgd) (See supplemental S4 File for list of genes and hgd implementation).

**Fig.4.**
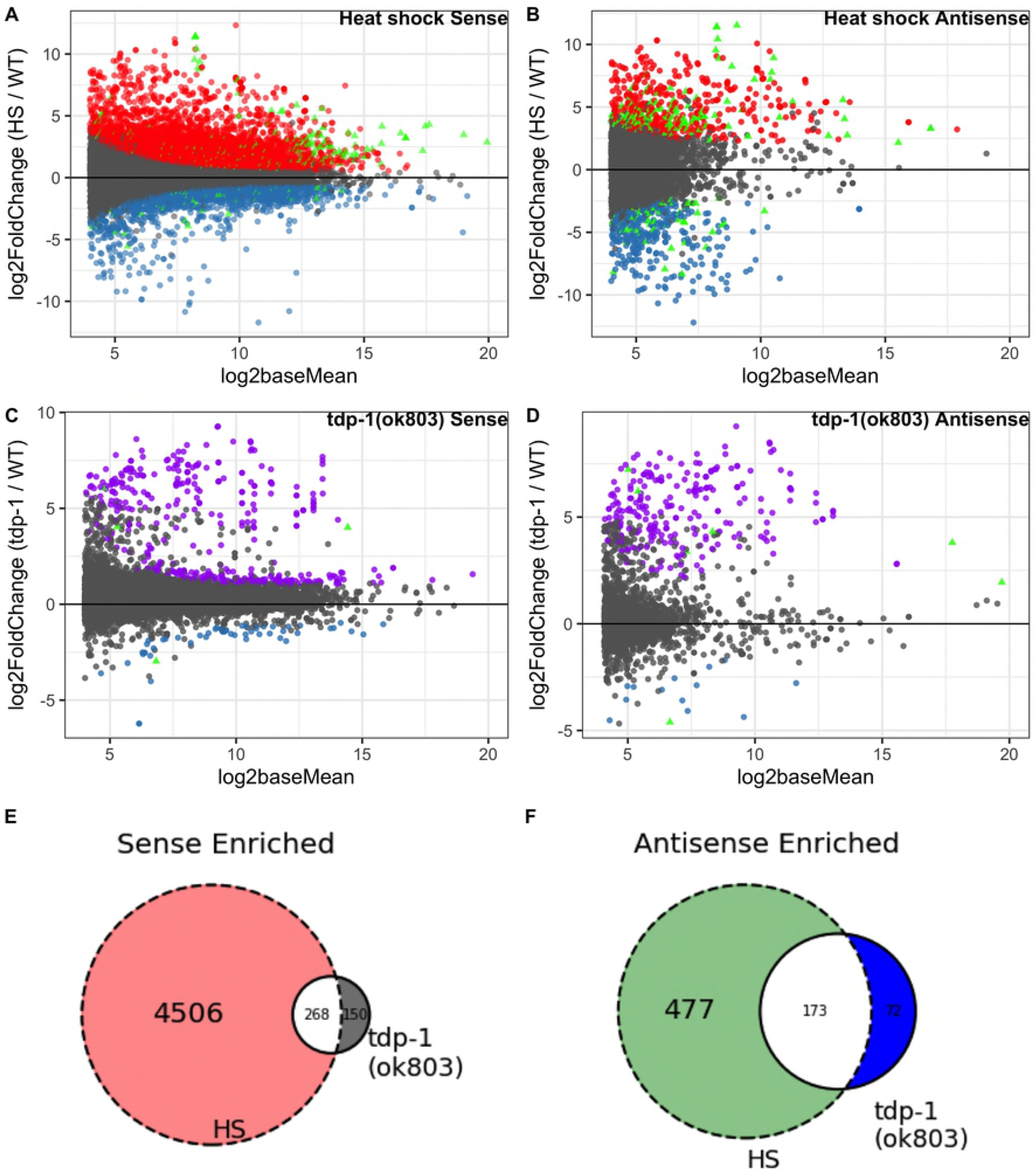
Comparison of J2 enriched sense and antisense transcripts in heat shock and *tdp-1(ok803)* worms. MA plots of significant (FDR <0.05) dsRNA enrichment for sense and antisense transcripts (analyzed independently) along with Venn diagrams of enrichment for enriched sense and antisense. (A) Heat shock over wild type J2 enriched sense transcripts with 4774 enriched (red) and 1669 depleted (blue). (B) Heat shock over wild type J2 enriched antisense transcripts with 650 enriched (red) and 477 depleted (blue). (A-B) enriched (180) and depleted (48) heat shock vs wild type transcripts found in both sense and antisense (green triangles). C: *tdp-1(ok803)* over wild type significant J2 enriched sense transcripts with 418 enriched (purple) and 59 depleted (blue). (D) *tdp-1(ok803)* over wild type significant J2 enriched antisense transcripts with 245 enriched (purple) and 14 depleted (blue). (C-D) enriched (6) and depleted (1) *tdp-1(ok803)* vs wild type transcripts found in both sense and antisense (green triangles). (E) Overlap of genes with significantly J2 enriched sense transcripts in both conditions compared to wild type worms. (F) Overlap of genes with significantly J2 enriched antisense transcripts in both conditions compared to wild type worms.

We used gene ontology (GO) analysis to investigate if transcripts enriched in the J2 immunoprecipitation fell into any functional categories. Using GOATOOLS [32], we found that many GO terms related to translation were significantly enriched (FDR < 0.05) in both the heat shock and *tdp-1(ok803)* J2-IP pools. In order to get the total number of genes involved in translation, which we call “translation related genes”, we combined lists of genes from all significantly enriched GO terms containing the words “translation”, “ribosome”, and “ribosomal”, then removed duplicate genes that were members of multiple GO terms (S3 File for methods). In sense J2 enriched transcripts, we find 234 translation related genes with heat shock, 27 translation related genes in *tdp-1(ok803),* and 19 translation related genes in the overlap. In the J2 depleted sense pool, heat shock contained 30 translation related genes, *tdp-1(ok803)*worms had no translation related genes, as well as no translation related genes in the overlap. In J2-enriched antisense transcripts, only heat shocked worms had 33 translation related genes. There was no translation related genes found in J2-depleted antisense transcripts (S5 File for list of significant genes and translation related genes, S6 File for GOATOOLS output).

### Enrichment of transcripts downstream of genes in the J2 pool

While examining the transcription of known heat shock-inducible genes, we noted in heat shocked populations an accumulation of read-through transcripts downstream of annotated genes (see example in Fig 5). Interestingly, some of these downstream-of-gene transcripts (DoGs) were also highly enriched in the J2-IP pool. While previous work has characterized the accumulation of downstream-of-gene transcripts in heat shocked cells from human [5] and mice [16], the phenomena has not previously been associated with dsRNA accumulation. To assay and find differential expression of these read-through regions across the whole genome, we created an algorithm called Dogcatcher (See methods and materials as well as GitHub README for algorithm explanation).

**Fig.5.**
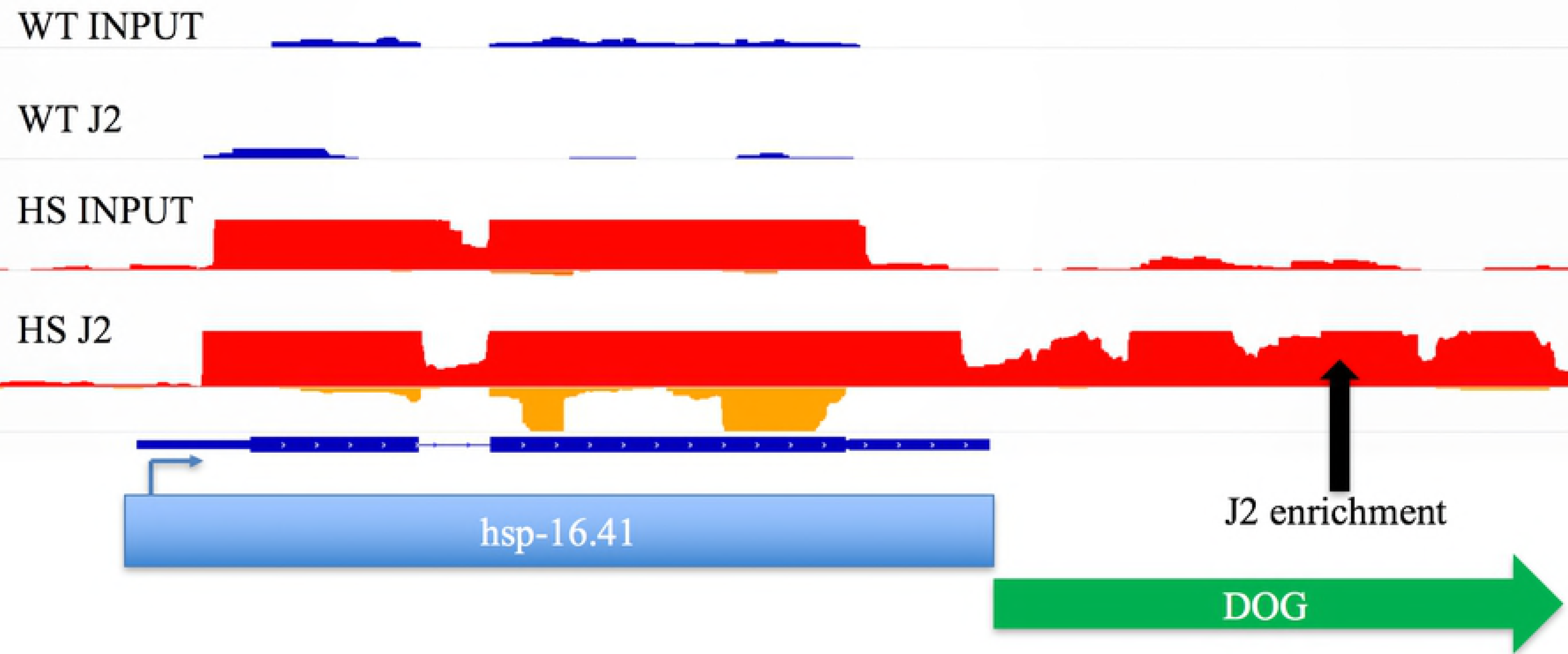
Aberrant transcription past the end of heat shock family genes showed enrichment in heat shock J2. Ribosomal subtracted ratio normalized histogram from the Integrative Genomics Viewer (IGV). On each track, the sense strand is on the top part of the histogram and antisense is on the bottom. Wild type (WT) sense (dark blue) and antisense (light blue), heat shock (HS) sense (red) and antisense (orange). Downstream of gene transcription (DoG) continues past the 3’ end of gene (green arrow).

Using the Dogcatcher algorithm, we were able to quantify downstream-of-gene transcripts in the J2-IP pool that can be missed using the standard *C. elegans* genome annotation. After heat shock, more read-through sections were significantly increased in the J2-IP pool than decreased (84 vs. 25). (Fig 6A). We found that for DoGs enriched in the J2-IP pool after heat shock, the majority correspond to protein coding genes (60%), followed by ncRNA (20%), pseudogenes (9%), and snoRNA (9%). We found far fewer significantly J2 enriched DoGs from *tdp-1(ok803)* (3) with no regions being depleted (Fig 6B). Interestingly, 2 out of the 3 DoGs in *tdp-1(ok803)* were also enriched in the heat shock J2 pool (S4 File for list DoGs and hgd implementation). From the significantly enriched GO terms of DoGs in heat shock and *tdp-1(ok803)* worms, only heat shocked worms had any significantly enriched GO terms which primarily consisted of histone genes (S7 File for list of DoGs, S8 File for GOATOOLS output). As a possible explanation for the formation of dsRNA at downstream of gene regions, we found DoGs to be enriched in terminal repeat sequences compared to a random intergenic downstream background. (S3 Fig, S3 File for methods).

**Fig.6.**
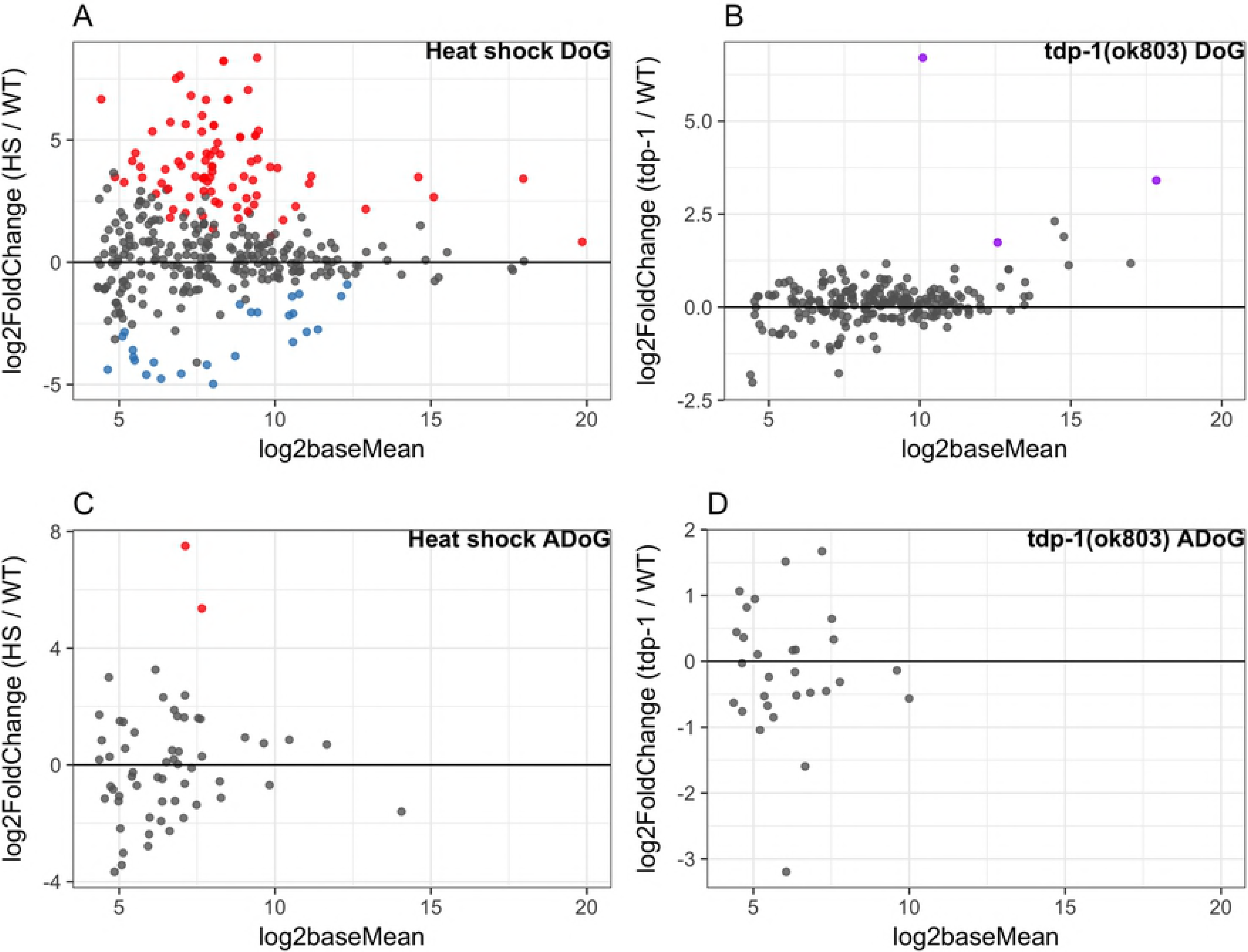
J2 enrichment of DoGs and ADoGs in heat shock and *tdp-1(ok803)* worms. MA plots of significant (FDR <0.05) dsRNA enrichment for DoGs and ADoGs. Non-significant genes added in with DoGs/ADoGs for DESeq2 normalization were taken out of the plots for clarity. (A) heat shock over wild type J2 enriched read-through sense transcripts with 84 enriched (red) and 25 depleted (blue). (B) *tdp-1(ok803)* over wild type significant J2 enriched read-through sense transcripts with 3 enriched (purple). (C) heat shock over wild type J2 enriched read-through antisense transcripts with 2 enriched (red). (D) No significant *tdp-1(ok803)* over wild type J2 enriched read-through antisense transcripts.

### Additional non-annotated transcripts are minimally enriched in the J2 pool after heat shock or *tdp-1* deletion

Next, we were curious if other sections around genes would show aberrant transcription in heat shock or *tdp-1(ok803)* worms. Expanding on the DoG nomenclature, the terms we use for the three other types of transcription flanking an annotated gene are as follows. Regions downstream of genes with antisense reads (ADoGs), sense reads in regions upstream/previous of the gene (POGs), and antisense reads in regions upstream/previous of the gene (APOGs) (See S4 Fig for visualizations and additional explanation). Importantly, novel areas of intergenic transcription are obtained by filtering out POGs that overlap with DoGs on the same strand, as well as ADoGs/APOGs that overlap DoGs or genes on the opposite strand (See S5 Fig for visualization of filtering). We did not find any significantly J2 enriched POGs or APOGs in either condition compared to wild type. We found a small amount of significant J2 enrichment in heat shock ADoGs (2) (Fig 6C) and no ADoGs enriched in *tdp-1(ok803)* (Fig 6D).

### Increased antisense transcription over genes associated with DoGS and ADoGS

Next, we were curious if any aberrant read-through transcription might overlap genes and contribute to increased antisense reads within the gene. We define an overlapped gene as any gene that has an ADoG associated with it or an opposite strand DoG with any overlap to the gene. We next define a significant overlapped gene as any gene that has significant antisense transcription as well as a significant ADoG or significant DoG that overlaps it on the opposite strand. From our significantly overlapped genes, we found 17 enriched and 5 depleted with heat shock, and only 4 enriched and no depleted in *tdp-1(ok803)* worms. We did not find any overlapped genes that were significantly enriched for GO terms related to translation (S5 File for list of overlapped genes, S3 File for overlap methods).

### Antisense read-through into *eif-3.B* in heat shocked worms

Visual inspection of DoG transcripts identified one transcript downstream of the ncRNA *W01D2.8 (doW01D2.8)* that ran into the gene *eif-3.B* on the opposite strand. (Fig 7A). *eif-3.B* is an ortholog of human EIF-3.B (eukaryotic translation initiation factor 3 subunit B) and is involved in translation initiation. As the *doW01D2.8* transcript was strongly increased by heat shock in both the total and J2-IP pools, we chose to target this transcript to confirm our RNA-seq data. Fluorescent *in situ* hybridization (FISH) was used as this could both demonstrate the accumulation of the *doW01D2.8* transcript and determine its cellular and subcellular (i.e., possible co-localization with J2 foci) distribution.

**Fig.7.**
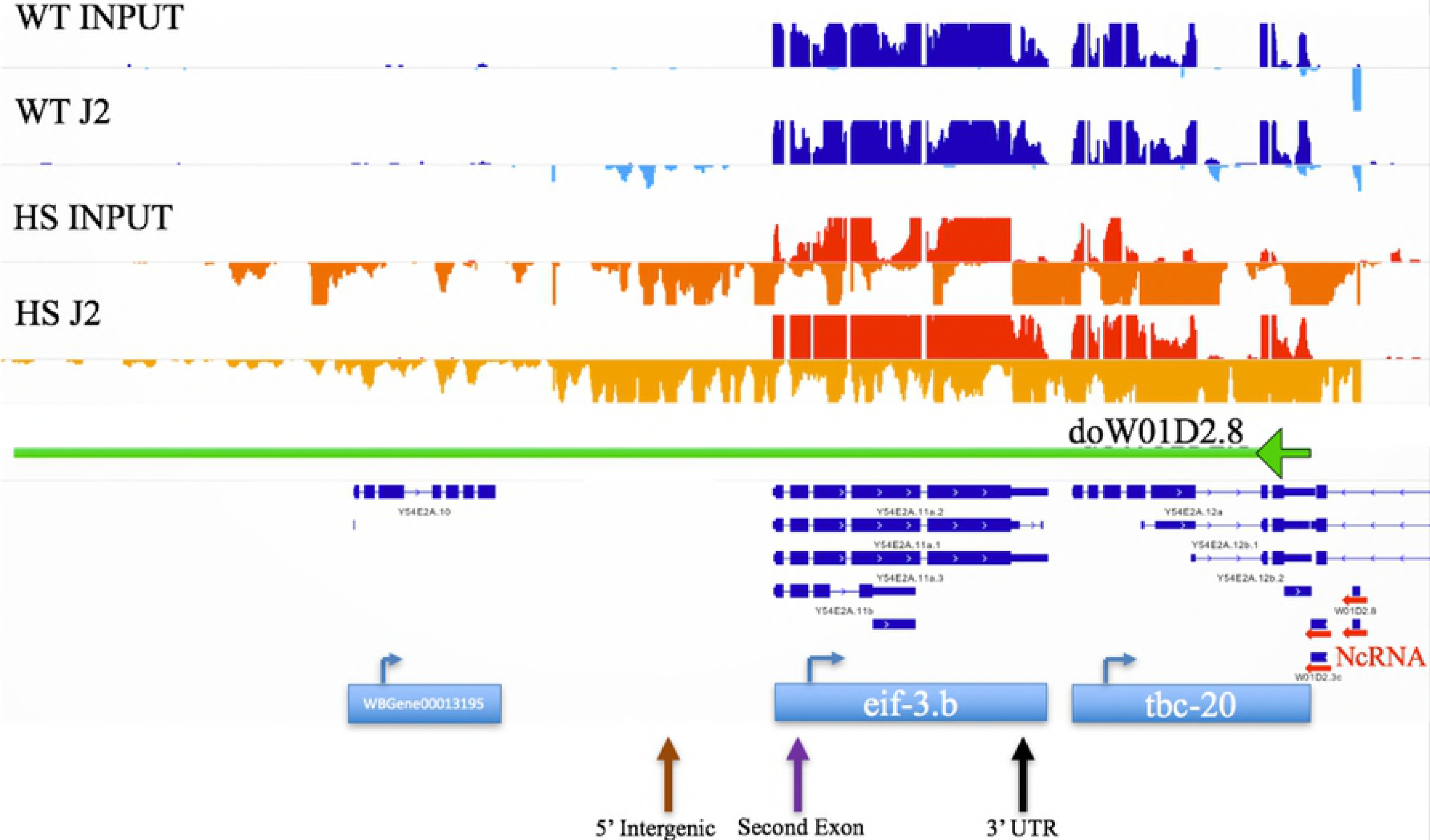
Heat shock induces transcripts antisense to the *eif-3B* locus. IGV view of *eif-3.B.* Tracks normalized by the ribosomal subtraction ratio, the sense strand is on the top part of the histogram and antisense is on the bottom. wild type (WT) sense (dark blue), WT antisense (light blue), heat shock sense (red), heat shock antisense (orange). Horizontal red arrows on the right show a cluster of ncRNAs including *W01D2.8* and transcription downstream of *W01D2.8 (doW01D2.8)*into *eif-3.B* (green arrow going left). Arrows on the bottom correspond to locations of primers for RT-PCR (brown: 5’ Intergenic, purple: Second exon, black: 3’ UTR).

Three strand-specific fish probes at the 5’ Intergenic region (5’ INT) (antisense), first 3 exons (sense), and the last exon along with the 3’ UTR (LE 3’UTR) (antisense) of *eif-3.B* (Fig 8D) were designed (list of probes in S1 File). First, we performed immunohistochemistry with the J2 antibody along with FISH for antisense transcripts that contain the last exon and 3’ UTR (see Material and Methods) (Fig 8A). We find that *doW01D2.8* is transcribed in this region with heat shock and commonly forms two foci per nucleus, but does not co-localize with dsRNA foci. The 5’ and 3’ doW01D2.8 probes do strongly colocalize in the nuclear foci (Fig 8B), consistent with a single transcript spanning this region. Additionally, when probing for the 5’ intergenic region (antisense) and first three exons (sense), we did not find the sense probes colocalizing to the antisense foci (S6 Fig). To inquire if the *eif-3B* antisense foci were a general site of transcript accumulation, we probed for *C30E1.9,* a long ncRNA that is highly expressed, forms nuclear foci, but is not induced in heat shock. We observed that this transcript does not overlap with the *eif-3B* antisense foci (Fig 8C). Lastly, we wanted to see if deletion of *tdp-1,* which does not lead to accumulation of *eif-3B* antisense transcripts, would alter heat shock induced accumulation of these transcripts. We found that the *tdp-1* deletion did not alter the formation of *eif-3B* antisense transcripts (Fig 8E).

**Fig.8.**
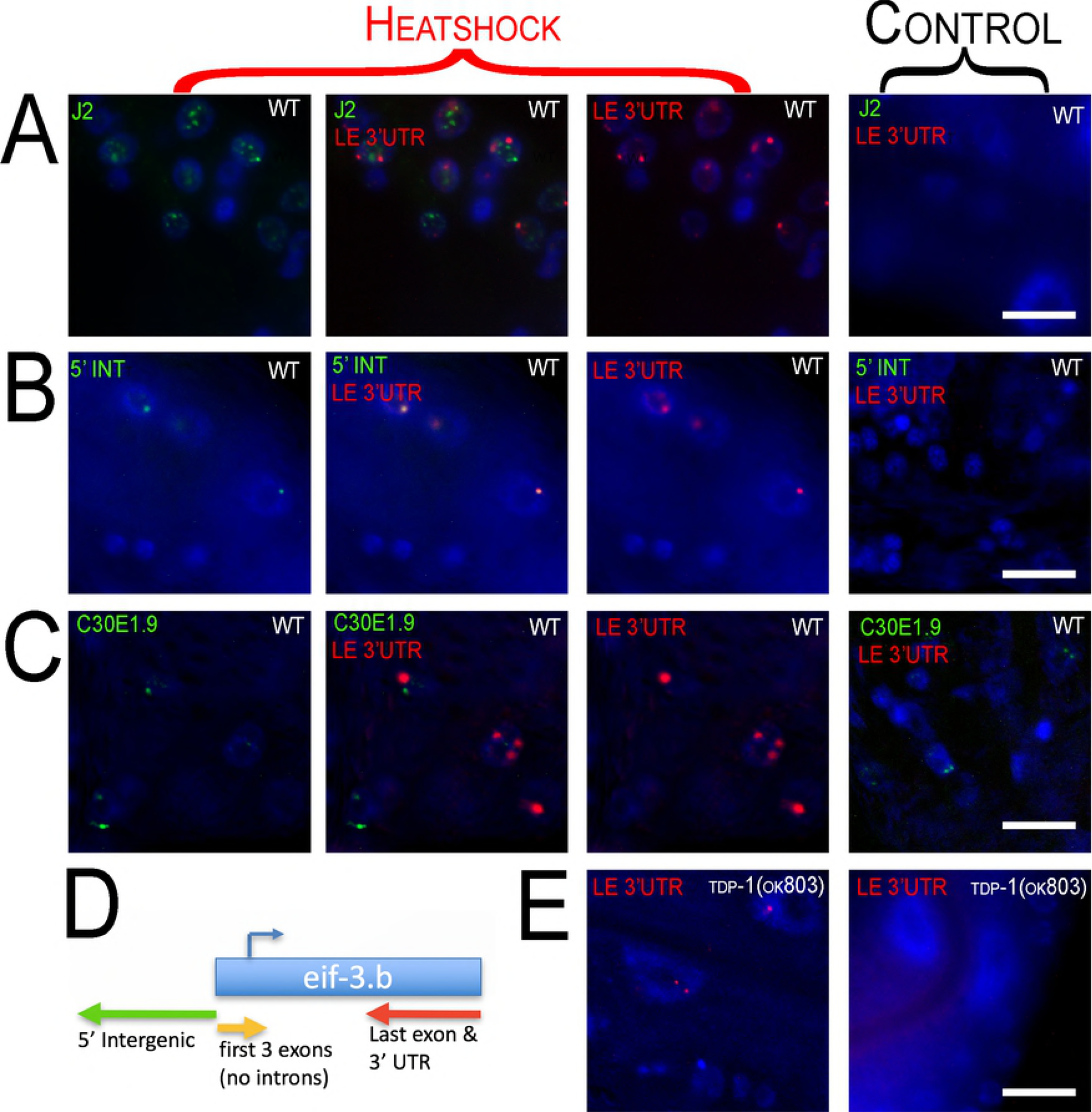
*Fluorescence in situ Hybridization (FISH)* of *eif-3.B regions.* 100x oil immersion images of worm hypodermal and neuronal cells. Heat shock panels are in the three columns to the left (merged channel in the middle column). Control panels show exposure from every channel (right column). Row (A) Immunohistochemistry with J2 antibody (green) along with FISH of *doW01D2.8* antisense to the last exon and 3’ UTR (LE 3’ UTR) (red) of *eif-3.B.* dsRNA and the antisense LE 3’UTR transcript aggregate into nuclear foci with heat shock and do not appear to co-localize. Row (B) FISH of *doW01D2.8* in two regions antisense to the 5’ intergenic region (5’ INT) (green) and last exon and 3’ UTR (LE 3’UTR) (red) of *eif-3.B.* Row (C) FISH of *doW01D2.8* antisense to the last exon and 3’ UTR (LE 3’UTR) (red) of *eif-3.B* and sense probe of ncRNA C3DE1.9 (green). Probing of C3DE1.9 is not affected by heat shock and C3DE1.9 is not induced by heat shock. C3DE1.9 and LE 3’UTR show no overlap. (D) Diagram of *eif-3.b* gene with FISH probe locations and orientation. (E) Heat shock of *tdp-1(ok803)* induces nuclear foci from probes antisense to the last exon and 3’ UTR (LE 3’UTR) of eif-3.B (left panel) and is not visible with no heat shock (right).

## Discussion

In our previous study [21] we established that in *C. elegans* deletion of *tdp-1* induces nuclear dsRNA foci. Here, we show that heat shock also induces nuclear dsRNA foci that partially overlap with HSF-1 nuclear stress granules. After heat shock, we find a general increase in the amount of dsRNA and expression levels of transcripts with dsRNA structure, assayed using the dsRNA-specific monoclonal antibody J2. In addition, we find that heat shock induces accumulation of antisense transcripts as well as novel downstream of gene transcripts. To our knowledge this is the first time heat shock has been shown to lead to the accumulation of these abnormal transcripts in an *in vivo* model.

dsRNA can form intra-or inter-strand base-pairing. Our data suggest that both types of dsRNA may be contributing to the dsRNA pool induced by heat shock. We find that novel downstream-of-gene transcripts are enriched in the J2-IP pool. These novel transcripts are enriched in inverted repeat sequences, which may be contributing to the formation of intra-strand dsRNA. Downstream-of gene transcripts also have the potential to generate transcripts antisense to neighboring genes on the other strand. This has been reported in the heat shock study by Vilborg et al [16], and we have noted similar examples in our data (see Figure 8). Using our new Dogcatcher algorithm, we have also documented novel transcripts originating in intergenic regions, which also have the potential to generate antisense transcripts. Indeed, we observe a general increase in antisense transcription after heat shock (see Figure 3), and antisense transcripts are enriched in the J2-IP, supporting the formation of inter-strand dsRNA. We note that the J2 antibody immunoprecipitation protocol used in our study will recover transcripts that have only partial (at least 40 nucleotides) dsRNA structure, thus it is feasible that some transcriptional regions we recover after J2-IP are single-stranded extensions of double stranded regions.

The accumulation of dsRNA transcripts after heat shock could be the result of altered RNA production and/or changes in RNA stability or turnover. Further studies (e.g. Pro-seq analysis) will be required to definitively determine the relative contribution of these cellular processes. Published studies demonstrate that loci susceptible to heat shock-induced downstream-of-gene transcription are marked by open chromatin before heat shock [16] and are depleted of the transcriptional termination factor CPSF-73 after heat shock [33]. These results suggest that altered transcriptional processing itself leads to the altered transcript accumulation after heat shock. However, the significant overlap of transcripts enriched in the J2 pool resulting from heat shock and from deletion of the *tdp-1* gene suggest that changes in RNA stability may be also contributing to transcript accumulation. TDP-1 is orthologous to mammalian TDP-43, and we have previously shown that human TDP-43 can act as an RNA chaperone in an *in vitro*assay [21]. Conceivably, heat shock could inhibit the function of TDP-1 or other similar RNA-binding proteins, leading to the formation of more dsRNA structure in existing transcripts.

We employed fluorescence in situ hybridization (FISH) to confirm heat shock-induced expression of DoG and antisense transcripts in the *eif-3.B* region, and to examine their subcellular localization. These novel transcripts were found in nuclear foci that did not overlap with either the HSF-1::GFP foci or the J2 dsRNA foci, and were typically limited to two spots in each nuclei. This two foci/nucleus distribution is very similar to the FISH characterization of DoG transcripts described by Vilborg et al, and strongly suggest that the *eif-3.B* loci transcripts are associated in *cis* with their site of production. These antisense transcripts clearly did not contribute to the foci detected by J2 immunostaining, and may reflect a general dysregulation of transcription at the *eif-3.B* locus. Identification of the dsRNA species present in the J2 foci induced by heat shock may require development of a protocol to purify these RNA granules, as we have identified thousands of transcripts enriched in the J2 pool, and have no additional insight as to which ones might be found specifically in the J2 foci.

A critical issue is whether the accumulation of novel transcripts and dsRNA after heat shock have a biological function. By characterizing transcriptional changes induced by a variety of stresses, Vilborg et al concluded that transcriptional read-through was not a random failure, and suggested it might have a functional role in stress responses. We have characterized the accumulation of dsRNA after heat shock, and by gene ontology analysis find that the sense and antisense transcripts in this pool (as well as the J2-IP pool in *tdp-1* deletion mutants) are enriched in genes involved in translation. Given that we find significant J2-IP enrichment of both sense and antisense transcripts from genes related to translation, it is tempting to speculate that the formation of inter-strand dsRNA might reduce the translation of these “translation-related transcripts”, leading to a down-regulation of global translation, a protective event against most cellular stress insults including heat shock. While we have no direct evidence that dsRNA-dependent translational downregulation happens after heat shock in *C. elegans,* we note that deletion of *tdp-1* has been reported to protect against proteotoxicity and increase lifespan [34]. Translational downregulation would presumably be protective against proteotoxicity, and postdevelopmental knockdown of translation initiation factors strongly increases lifespan in *C. elegans* [35].

## Acknowledgements

We would like to thank Anna Vilborg for supplying the perl scripts from her initial DoG publication [5]. Some nematode strains were provided by the Caenorhabditis Genetics Center, funded by the NIH National Center for Research Resources.

## Supporting information

S1 Fig. Comparison of heat shock J2 enriched transcripts significant in both sense and antisense.

S1 Table. Worm strains used in this study.

S1 File. Probes used in Fluorescence in situ Hybridization (FISH) of eif-3.B regions.

S2 Fig. Comparison of *tdp-1(ok803)* J2 enriched transcripts significant in both sense and antisense.

S2 Table. Quantification of occurrence of HSF-1 and J2 foci over time (Raw data).

S3 Fig. Number of Terminal Inverted Repeats (TIR) overlapping downstream regions.

S2 File. List and sequences of adapters used in trimming.

S3 File. Bioinformatic methods.

S4 Fig. Dogcatcher flattening and nomenclature.

S4 File. Hypergeometric Distribution and list of genes/DoGs used in calculation for Heat shock and *tdp-1(ok803).*

S5 File. List of significant genes, translation associated genes, and overlapped genes.

S5 Fig. Dogcatcher additional filtering.

S6 File. Significantly enriched GO terms for Heat shock and *tdp-1(ok803)* genes.

S6 Fig. Sense and antisense *eif-3B* transcripts do not colocalize.

S7 File. List of significant DoGs, ADoGs, PoGs, APoGs.

S8 File. Significantly enriched GO terms for Heat shock DoGs.

